# Meta-analysis of the black soldier fly (*Hermetia illucens*) microbiota based on 16S rRNA gene amplicon sequencing

**DOI:** 10.1101/2022.01.17.476578

**Authors:** Freek IJdema, Jeroen De Smet, Sam Crauwels, Bart Lievens, Leen Van Campenhout

**Affiliations:** CLMT Research Group for Insect Production and Processing, Department of Microbial and Molecular Systems (M^2^S), KU Leuven, Campus Geel, Geel, Belgium; KU Leuven, Leuven Food Science and Nutrition Research Centre (LFoRCe), Leuven, Belgium; CMPG Laboratory for Process Microbial Ecology and Bioinspirational Management (PME&BIM), Department of Microbial and Molecular Systems (M^2^S), KU Leuven, Leuven, Belgium; Leuven Plant Institute (LPI), KU Leuven, Leuven, Belgium

**Author notes:** Corresponding author: CLMT Research Group for Insect Production and Processing, Department of Microbial and Molecular Systems (M^2^S), KU Leuven, Kleinhoefstraat 4, B-2440 Geel, Belgium.

**Keywords:** 16S rRNA gene, bacteria, Black Soldier Fly, Hermetia illucens, meta-analysis, microbiota

## Abstract

Black soldier fly larvae (BSFL) belong to the most widely reared insect species as an alternative protein source at industrial scale. Bacteria in the larval gut can provide benefits for the animal, though some bacteria can also be pathogenic for the insect. Accurate characterization of the BSFL microbiota is important for the production of BSFL in terms of yield and microbiological safety. In this study, 16S ribosomal RNA gene sequence data sets from 11 studies were re-analysed to gain better insights in the BSFL gut microbiota, potential factors that influence their composition, and differences between the gut and the whole larvae microbiota. A core gut microbiota was found consisting of members of *Enterococcus, Klebsiella, Morganella, Providencia*, and *Scrofimicrobium.* Further, the factors “Study”, “Age” and “Feed” significantly affected the microbiota gut composition. When compared to whole larvae, a significantly lower number of observed zero-radius Operational Taxonomic Units and a lower diversity was found for gut samples, suggesting that the larvae harboured additional microbes on their cuticle or in the insect body. Universal choices in insect sample type, primer selection and bio-informatics can strengthen future meta-analyses and improve our understanding of the BSFL gut microbiota towards the optimization of insect production.

## Introduction

Insects are one of the most promising alternative sources to sustainably produce proteins. They not only contain high quality protein and essential nutrients, as they also have a high potential for efficient upgrading of waste streams. In this regard, Black Soldier Fly larvae (BSFL) (*Hermetia illucens* L., Diptera: Stratiomyidae) are one of the most promising insect species to be used as an alternative protein source in animal feeds. The species, originally traced to the Americas, is present in most tropical and temperate regions of the world (Sheppard *et al.*, 1994). Although their nutritional value varies, depending on their rearing substrate, crude proteins of BSFL can account for 32-45% of the total dry weight of the larvae, while fat constitutes about 15-49% of their total dry weight. In addition, their amino acid profile is similar to that of fishmeal (Belghit *et al.*, 2019; Caligiani *et al.*, 2018; Diener *et al.*, 2009; Henry *et al.*, 2015; Makkar *et al.*, 2014). Though naturally occurring in chicken, pig and cow manure, BSFL have been successfully reared on diverse organic waste streams and are being cultivated at industrial scale (Miranda *et al.*, 2020; Tomberlin & Van Huis, 2020; Yang *et al.*, 2020). Altogether, this makes BSFL a highly suitable source for traditional animal protein.

Microbes in the gut can substantially support nutrient uptake processes and the host immune system (O’Hara *et al.*, 2020). The gut microbiota has therefore been an important subject of study in traditional animal health and production (Holman & Gzyl, 2019; O’Hara *et al.*, 2020) and likewise, it is useful to investigate the microbiota in insect production. Several studies have investigated the microbiota in and on BSFL, aiming to get a better understanding of the BSFL microbial community composition and function to aid their industrial production, as well as to assess potential microbial hazards associated with their consumption (Khamis *et al.*, 2020, Wynants *et al.*, 2018). The dissected insect gut is often used for microbiota analysis, although some studies use the whole larvae as their sample material. The choice for a specific sample type can depend on the focus of the study. However, there might be a difference between the microbiota composition of both sample types. Aside from the insect gut, microorganisms can be present in the insect haemolymph (Aronson *et al.*, 1986) or on the surface of the insect (Ulanova *et al.*, 2020). External bacteria could enter the larvae through oral ingestion but also through wounds or opening in the insect cuticula (Joosten *et al.*, 2020). It is therefore important not only to consider the gut microbiota, but also to take into account microorganisms colonizing other locations in the insect.

Whereas bacterial communities have been traditionally studied through culture-based methods, focus has shifted towards next-generation sequencing approaches that target a taxonomic marker such as the ribosomal RNA (rRNA) gene. Compared to culture-dependent methods, sequencing-based approaches have several advantages (Hiergeist *et al.*, 2015). It is well known that not all microorganisms can be cultivated in standard laboratory conditions. Furthermore, sometimes important microbial groups are missed due to culture-dependent biases, such as the appearance of microorganisms in a viable but non-culturable state and/or the possibility that isolation media favor the cultivation of specific microorganisms, that outcompete others. Additionally, microorganisms may remain undetected when they occur at cell densities below the limit of detection of the plate methods. These drawbacks can be circumvented by next-generation sequencing technologies, enabling an in-depth characterization of microbial communities without culturing (Oulas *et al.*, 2015).

In general, 16 phyla of bacteria have been identified in the BSFL gut, with Firmicutes, Bacteroidetes and Proteobacteria being the most dominant phyla (Huang *et al.*, 2020; Zhan *et al.*, 2020; Zhang X. *et al.*, 2020). Ao *et al.* (2020) found that the amount of total nitrogen and fat was positively correlated with the relative abundance of *Providencia* in the gut of BSFL fed on chicken and swine manure. This suggests that *Providencia* might play an important role in protein and lipid conversion in the gut (Ao *et al.*, 2020). *Providencia* was also found at high relative abundance in other studies focussing on BSFL microbiota (Gorrens *et al.*, 2021; Raimondi *et al.*, 2020; Wynants *et al.*, 2018; Zheng *et al.*, 2013). Additionally, Bruno *et al.* (2019) suggested that members of the genera *Sphingobacterium* and *Dysgonomonas* may play a role in the degradation of polysaccharides in the BSFL gut. In contrast, there are also bacteria in the BSFL gut which may have a deleterious effect on their host. For example, Wu *et al.* (2020) found members of bacterial families in the BSFL gut that may be detrimental, including the families *Brucellaceae* and *Enterobacteriaceae*, and the genus *Campylobacter.* Zheng *et al.* (2013) found potential pathogenic *Enterobacteriaceae* members in all life stages of BSFL raised on chicken feed. Further, it has been found that environmental factors like rearing temperature and diet have a strong impact on the BSFL microbial community composition and diversity. Raimondi *et al.* (2020) found a negative correlation between temperature and the relative abundance of *Providencia*, but a positive correlation with several other bacteria (*e.g. Alcaligenes, Pseudogracilibacillus, Proteus).* For many insect species it has been shown that diet has a large impact on the gut microbiota (Engel & Moran, 2013). This has also been demonstrated for BSFL (Ao *et al.*, 2020; Bruno *et al.*, 2019; Osimani *et al.*, 2021). Furthermore, studies have suggested that geographic origin and the inhouse bacterial community of rearing facilities may influence the microbiota composition of edible insects (Khamis *et al.*, 2020; Wynants *et al.*, 2018).

Although these studies have greatly improved our knowledge on BSFL microbiota and potential factors determining the microbial community composition, it remains challenging to identify general trends and draw general conclusions over the different studies performed. Studies have been performed using different PCR primers and regions, which may affect the outcome of microbiota studies (Albertsen *et al.*, 2015; Fouhy *et al.*, 2016; Rintala *et al.*, 2017; Tremblay *et al.*, 2015; Zhang *et al.*, 2018). Additionally, different bioinformatics data analysis pipelines have been used to analyse the data (*e.g.* MOTHUR, QIIME, USEARCH) with various workflows. Moreover, there is a growing tendency to replace the use of operational taxonomic units (OTUs) based on a sequence similarity cut-off by the analysis of exact sequence variants (also termed zero-radius OTUs (zOTUs)) or amplicon sequence variants (ASVs) (Callahan *et al.*, 2017; Edgar, 2016), increasing taxonomic resolution. Altogether, this makes it very difficult to compare different studies. Therefore, to increase our understanding of the BSFL microbiota and the underlying mechanisms shaping the microbial communities, a profound meta-analysis of the different studies performed is needed, in which all data is analysed in the same manner, *i.e.* using the same bioinformatic pipeline.

In this study, the BSFL gut microbiota is characterized in detail by processing raw 16S rRNA gene sequencing data from different publicly available studies through the same bioinformatic pipeline. A first objective was to investigate whether a specific set of gut bacteria can be identified despite varying experimental conditions (the “core” microbiota), and if so, which genera or species belong to the core gut microbiota. Secondly, we aimed to study which factors, such as type of feed and age of the larvae at harvest, shape the bacterial composition in BSFL gut samples. And finally, a comparison was made between the microbiota composition found in the gut of BSFL and in larvae as a whole to investigate differences between these two sample types.

## Materials and methods

### Data acquisition and quality filters

A total of 11 studies were included in the meta-analysis, and they are described in Table 1. The studies were retrieved through a literature search (performed in 2021) using the key words *“microbiome”, “black soldier fly”, “microbiota”* or *”Hermetia illucens”* in Scopus, PubMed and Google Scholar. Additionally, potential datasets were identified through a search in the Sequence Read Archive (SRA) database from NCBI PubMed (available from: http://www.ncbi.nlm.nih.gov/pubmed) by entering the key words *“black soldier fly”* or *“Hermeiia illucens”* (accessed in 2021), resulting in 40 and 41 BioProject hits, respectively. To be included in the meta-analysis, all studies were required (i) to have analysed the BSFL microbiota, (ii) to have analysed samples that reflect a ‘natural’, inartificial state of the microbiota composition, which excludes studies where bacterial strains were injected or introduced to the BSFL or its feed, (iii) to have used high-throughput amplicon sequencing of the 16S rRNA gene, (iv) to have used a sequence length of at least 250 bp, and (v) to have associated metadata and quality score files. BSFL studies that did not meet these criteria are summarized in Supplementary Table S1. For the 11 studies that met our criteria, the raw sequence data were collected in fastq-format using the NCBI Sequence Read Archive (SRA)-Toolkit (https://github.com/ncbi/sra-tools). Overall, the studies included BSFL gut samples originating from 13 countries worldwide and whole larvae samples from five European countries. All samples were collected between 2016 and 2021.

**Table 1.**
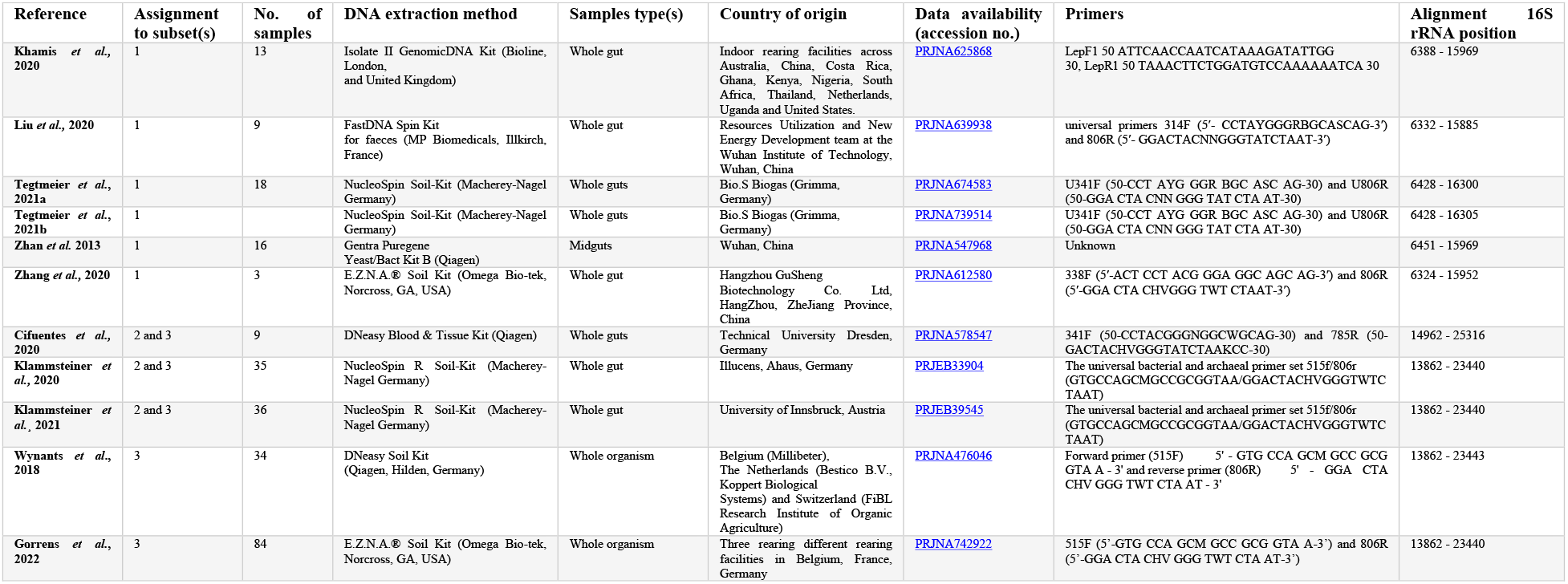
Overview of the raw data collected from each paper. Sample meta-information was collected from the papers as well as from the supplementary metadatafile included with the dataset when it was availble on the NCBI website. The sequencing platform used in these studies was Illumina Miseq. Not all papers included their primer use in either of these sources. Number of samples represent the fastq files of the larval samples of these studies that were used in this meta-analysis. The 16S rRNA region are determined through an alignment on the Da Silva Database (v123). Datasets were devided and analysis seperately: Subset 1 consisted of six datasets containing gut sample sequences located on the 6388-15969 region; Subset 2 consisted of three datasets containing gut sample sequences located on the 13862 – 25316 region; Subset 3 consisted of three datasets containing gut sample sequences and two dataset containing whole larvae sample sequences located at the 13862 – 25316 region.

### Processing of the 16S rRNA gene sequences

After aligning the obtained sequences against the Silva Living Tree Project v123 (LTP v 123) database, it became clear that six datasets aligned together (6332 – 16305), while the other five datasets aligned to a region more downstream in the 16S rRNA gene (13862 – 25316). As there was only very little overlap between both regions (less than 94 bp in 99% of the alignments), the overall data set was split in three subsets. Subset 1 consisted of six data sets of bacterial sequences from BSFL gut samples spanning the 6388 – 15969 region (corresponding to the V3-V4 region of the 16S rRNA gene). Subset 2 consisted of three data sets containing gut sample sequences covering the 13862 – 25316 region (*i.e.* the V4-V5 region), and Subset 3 consisted of three data sets containing sequences from gut samples and two sets with sequences from whole larvae samples located at the 13862 – 25316 region (V4-V5 region) (Yarza *et al.*, 2014) (Table 1). Further analysis of each of the three separate subsets is described below.

Obtained demultiplexed pair-end reads (with primer and barcodes removed) were merged using USEARCH (Edgar, 2010) fastq-mergepairs to form consensus sequences with no more than 10 mismatches allowed in the overlap region. Sequences were trimmed at a length of 250 bp with a maximum estimated error of 0.5 (VSEARCH-fastq_filter option (Rognes *et al.*, 2016)). Unique sequences were classified using the Silva Living Tree Project database v123 and non-bacterial sequences (*i.e.* chloroplasts, mitochondria, Archaea, Eukaryota) were removed using MOTHUR’s ‘classify.seqs()’ and ‘remove.lineage()’ commands (Schloss *et al.*, 2009). The resulting decontaminated sequences were classified into zOTUs by the UNOISE3 algorithm as implemented in USEARCH. zOTUs occurring at a relative abundance below a threshold of 0.1% per sample were set to zero. zOTUs were identified using the SINTAX algorithm implemented in USEARCH based on the Silva Living Tree Project v123 database. Further, the identity of the most important zOTUs was verified with a BLAST search in GenBank against exemplar species or “type materials” (Federhen, 2015). When no significant similarity was found with type materials (<97 % identity), the BLAST analysis was performed against entire GenBank. Identification was based on the highest max score, identity percentage and the lowest E-value.

### Data analysis and statistics

#### Core bacteria

First, the Subset 1 and Subset 2 data sets were used to define the “core microbiota” occurring in BSFL guts. Core microorganisms were defined in two ways, in one definition based on a rather stringent criterium of zOTUs being present in at least 80% of all samples and a second definition of zOTU presence in more than 50% of all samples (both definitions will be referred to later on). To this end, for each sample, zOTUs were scored as being present or absent. Presence or absence and relative abundance of the zOTUs over all samples were analysed using the *phyloseq* (McMurdie & Holmes, 2013) and *microbiome* (Lahti & Shetty, 2012-2019) packages in R 4.0.4 (R Core Team, 2021).

#### Factors determining the community composition

For the Subsets 1 and 2, beta diversity estimates (permutational analysis of variance (PERMANOVA) and visualized by non-metric multidimensional scaling (NMDS)) of Hellinger-transformed relative abundance data were used to study the effect of three experimental factors potentially influencing the gut microbiota composition of BSFL, including “Study”, “Age” and “Feed”. The factor “Study” refers to the fact that each study performed may vary in other experimental parameters that might influence the microbiota composition (*i.e.* variation in the strains of BSFL used, presence of inhouse bacterial genera, difference in scale at which larvae were reared, difference in DNA-extraction methods, and other possible experimental differences). For the factor “Age”, two subgroups were differentiated, including a subgroup “Young” and a subgroup “Old”. The subgroup “Young” consisted of larval samples harvested at an age of 4 – 14 days, or described as “early instar larvae” (stage L1 and L2). The subgroup “Old” was composed of larval samples harvested at an age of 15 days and above, or described as L3-L5 larvae or “prepupae”. Larvae indicated with an age of “14 – 17 days” were categorized as “Old”. With regard to the parameter “Feed”, samples were grouped into six categories based on the different food sources used. The categories were: “Agricultural sidestreams”, “Animal feed”, “Food/oil waste”, “Fruit/vegetable waste”, “Manure” and “Oily sidestreams”. Larval samples that were reared on substrates that would directly influence the microbiota (such as substrates supplemented with antibiotics or inoculated with microbial strains) were discarded from the analysis, as the microbiota of these samples possibly did not reflect the natural diversity in the samples. Also, six samples reared on poultry blood (Wynants *et al.*, 2018) were removed from the analysis, as this type of rearing substrate is not commonly used in BSFL production. Furthermore, 13 samples originating from the study of Khamis *et al.* (2020) did not provide any information about larval age or rearing substrate, and they were therefore also discarded from this part of the analysis.

#### Comparison between gut and whole larvae samples

Differences in the microbiota composition between gut and whole larvae samples (Subset 3) were evaluated using the alpha diversity indices zOTU richness, Shannon’s diversity index and Simpson’s diversity. The student’s unpaired t-test (in case of normality and equal variances) and the Wilcoxon Rank Sum test using the Bonferroni’s correction (in case of non-normality) were used to reveal any statistically significant differences between both subsets of samples. Further, differences between both subsets were analysed with beta diversity estimates using PERMANOVA and visualized by NMDS, based on a Bray-Curtis Matrix of Hellinger-transformed relative abundance data.

## Results

### Meta-analysis characteristics

Data collection resulted in sequences originating from 145 BSFL gut samples and 114 whole BSFL samples, located at different regions of the 16S rRNA gene. These sequences were divided into three subsets, as described earlier based on 16S rRNA gene region (Subset 1 and 2) or sample type (Subset 3). Subset 1 consisted of 7,666,062 sequences originating from 65 BSFL gut samples, Subset 2 consisted of 5,856,794 sequences from 80 BSFL gut samples, and Subset 3 consisted of 8,719,940 sequences representing the same 80 BSFL gut samples combined with 114 whole larvae samples. Subset 1 yielded a total of 6,652 zOTUs, of which 281 zOTUs accounted for 95% of the total number of reads. Subset 2 yielded a total of 1,736 zOTUs, of which 46 zOTUs accounted for 95% of the total number of reads. Subset 3 contained a total of 3,276 zOTUs, of which 154 zOTUs accounted for 95% of the total number of reads. Rarefaction curves for the samples in all three subsets were calculated and visualised in Supplementary Figure S1-S3. The saturation of these rarefaction curves indicates that the sequence depths of the samples were sufficient to cover microbial diversity.

### Identification of the core gut bacteria of BSFL

Core bacteria were identified by analysing both Subset 1 and Subset 2 sequences. Although the number of studies and samples differed between the two data sets, the evaluation of the alpha diversity revealed no significant differences in the number of observed zOTUs per sample, zOTU richness nor Shannon’s and Simpson’s diversity indices between Subsets 1 and 2 (Figure 1).

**Figure 1.**
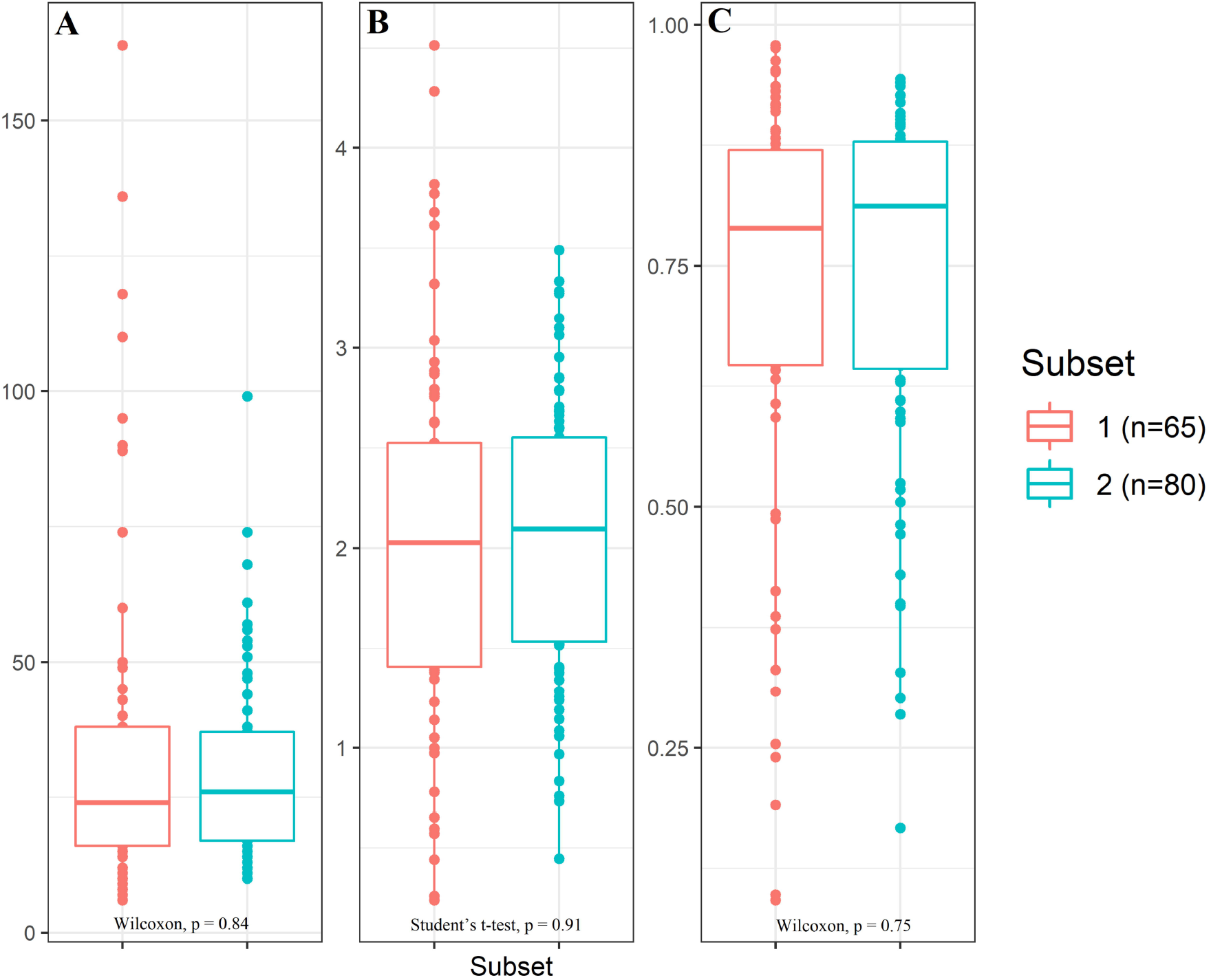
Alpha diversity measurements for BSFL gut samples Subsets 1 and 2. Observed zOTU richness (A), Shannon’s diversity index (B) and Simpson’s diversity (C) are visualized for BSFL gut samples from Subset 1 (red, n=65) and Subset 2 (blue, n = 80) using boxplots. Student’s unpaired t-tests and Wilcoxon Rank Sum tests are used for significance, p-values of these tests are provided under each boxplot.

An overview of the identification of the most prevalent zOTUs from Subsets 1 and 2 is provided in Supplementary Table S2. When considering the most prevalent bacteria in Subset 1, there were three zOTUs matching the strict definition of the core microbiota as being present in over 80% of the samples, representing members of *Enterococcus* (zOTU2, 87.7%), *Morganella* (zOTU1, 84.6%) and *Providencia* (zOTU9, 80.0%). When applying the less strict definition of the core microbiota as being present in at least 50% of the samples, six additional zOTUs were found, representing members of the genera *Proteus* (zOTU7, 73.8%), *Klebsiella* (zOTU13, 64.6%), *Enterococcus* (zOTU5, 64.6%; zOTU41, 69.2%), *Scrofimicrobium* (zOTU21, 56.9%) and *Enterocloster* (zOTU15, 52.3%) (Figure 2).

**Figure 2.**
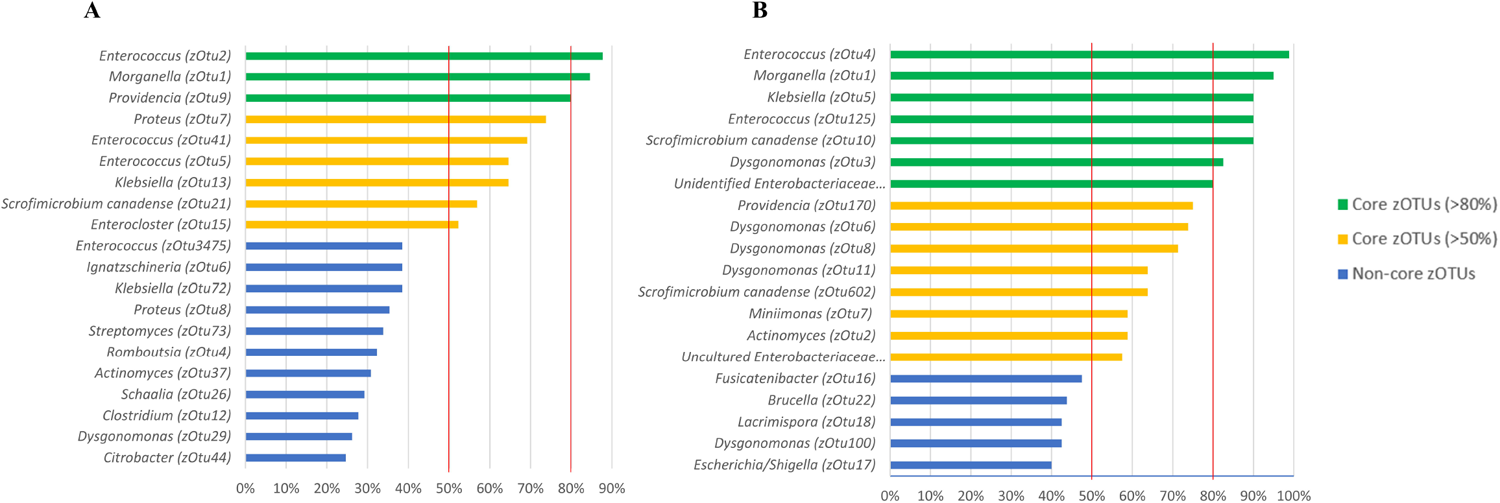
Most prevalent zOTUs in the gut samples from Subsets 1 (A) and 2 (B). Prevalence of zOTUs was based on absence and presence of zOTUs across all samples (n=65 for Subset 1, n = 80 for Subset 2) for each separate subset. These most prevalent zOTUs were identified using BLAST against type materials. When blasting of zOTUs yielded less than 97% identification, these sequences were blasted against the entire GenBank. zOTUs were considered to be part of the BSFL core microbiota when they were prevalent in over 80% (strict definition, green bars) and 50% (mild definition, yellow bars) of all samples within each group. Blue bars represent prevalent species under these threshold values.

In Subset 2, seven bacterial zOTUs were present in more than 80% of the samples investigated, representing members of *Enterococcus* (zOTU4, 98.8%; zOTU125, 90.0%), *Morganella* (zOTU1, 95.0%), *Scrofimicrobium* (zOTU10, 90.0%), *Klebsiella* (zOTU5, 90.0%), *Dysgonomonas* (zOTU3, 82.5%) and an unidentified member of the *Enterobacteriaceae* family (zOTU137, 80.0%). Eight additional zOTUs were included in the core microbiota when the prevalence threshold was lowered to 50% of all gut samples. These additional zOTUs corresponded to species of *Providencia* (zOTU170, 75.0%), *Dysgonomonas* (zOTU6, 73.8%; zOTU8, 71.3%; zOTU11, 63.8%), *Scrofimicrobium* (zOTU602, 63.8%), *Miniimonas* (zOTU7, 58.8%), *Actinomyces* (zOTU2, 58.8%) and an unknown *Enterobacteriaceae* genus (zOTU13, 57.5%) (Figure 2).

### Factors affecting the bacterial composition of BSFL gut

NMDS clustering of the data from Subsets 1 and 2 reveals clustering of samples based on age for both subsets (Figure 3A and 3D). Clustering based on “Feed” can be observed in Figures 3B and 3E, respectively, for Subset 1, and based on “Study” in Figures 3C and 3F, respectively, for Subset 2. PERMANOVA of the Hellinger transformed Bray-Curtis dissimilarities revealed that each of these three factors had a significant effect on the total bacterial composition found in the gut samples of both subsets, and when they were combined, they explained nearly 40% of all the variation (Table 2). In Subset 1, the factor “Feed” explained most of the variation between the samples, followed by “Study”. The factor “Age” only accounted for a small proportion of the variation among samples. For Subset 2, “Study” and “Feed” also accounted for a larger proportion of the variation relative to “Age”.

**Figure 3.**
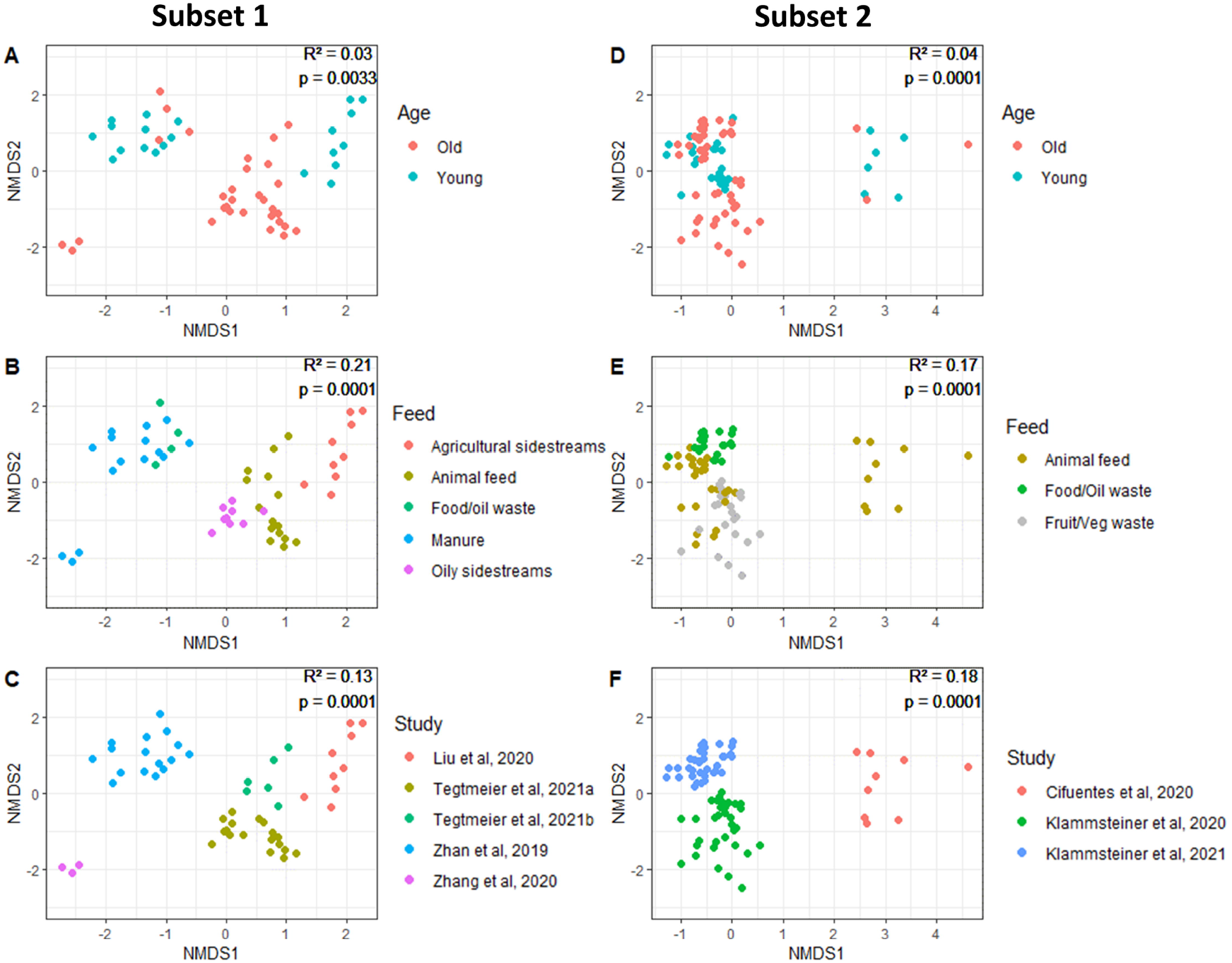
NMDS plots showing clustering of larval gut samples based on three factors, based on Bray-Curtis matrices of Hellinger-transformed relative abundance data. Scatterplots show clustering of gut samples of Subset 1 (A, B, C) and Subset 2 (D, E, F) for the three varying factors “Age”, “Feed” and “Study”. Results of an PERMANOVA analysis are shown in the upper right corner of each plot and reveal the significance (p-value) of each factor and the amount of variation that factor explains (R^2^).

**Table 2.** Results of PERMANOVA analysis of factors affecting the gut microbiota of BSFL. Tested factors had a significant effect on the microbiota composition found in gut samples of Subsets 1 and 2. Statistical testing of Bray-Curtis dissimilarities of Hellinger-transformed relative abundance data was done using PERMANOVA with the adonis function in R with 9,999 permutations.

|  | **Subset 1** |  | **Subset 2** |  |
| --- | --- | --- | --- | --- |
| **Factor** | **R<sup>2</sup>** | **P-value** | **R<sup>2</sup>** | **P-value** |
| *Age* | 0.03 | 0.0033 | 0.04 | 0.0001 |
| *Feed* | 0.21 | 0.0001 | 0.17 | 0.0001 |
| *Study* | 0.13 | 0.001 | 0.18 | 0.0001 |

### Difference in bacterial composition between gut and whole larval samples

Alpha diversity determinations of Subset 3 revealed that the observed number of zOTUs and bacterial diversity was significantly higher in the “Whole larvae” samples than in the “Gut” samples, as illustrated in Figure 4. A beta-analysis NMDS-plot of Bray-Curtis values (Figure 5) showed clear grouping based on the factor “Tissue” (representing either “gut samples” or “whole larvae samples”). Furthermore, PERMANOVA results showed that the factor “Tissue” had a significant (p = 0.0001) effect, although a relatively low share in the variation was explained by this factor (R^2^ = 0.05) (Table 3).

**Figure 4.**
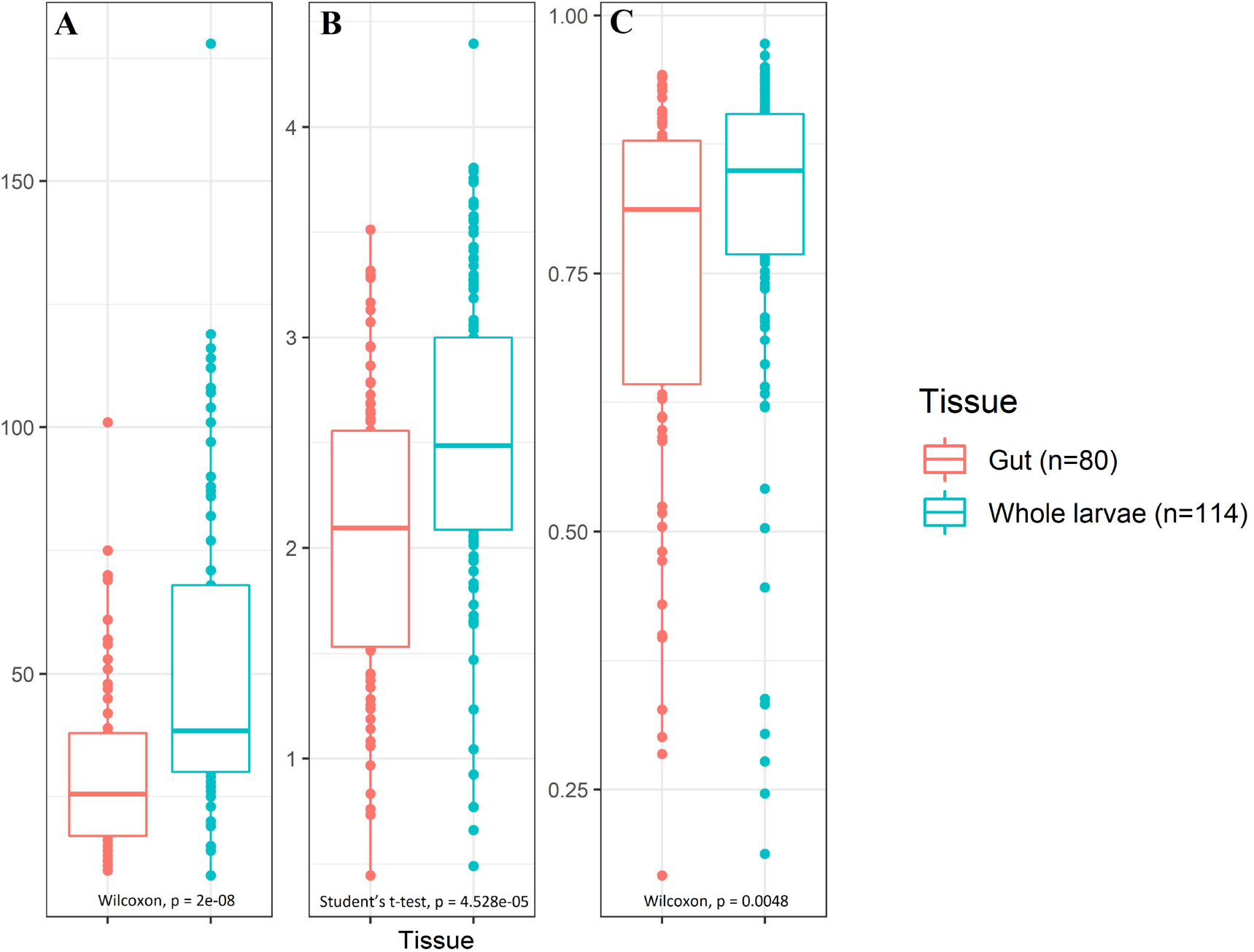
Alpha diversity measurements for gut and whole larvae samples from Subset 3. Observed zOTU richness (A), Shannon’s diversity index (B) and Simpson’s diversity (C) are visualized for gut (red, n=80) samples and whole larvae (blue, n = 114) samples using boxplots. Student’s unpaired t-tests and Wilcoxon Rank Sum tests are used for significance, p-values of these tests are provided under each boxplot.

**Figure 5.**
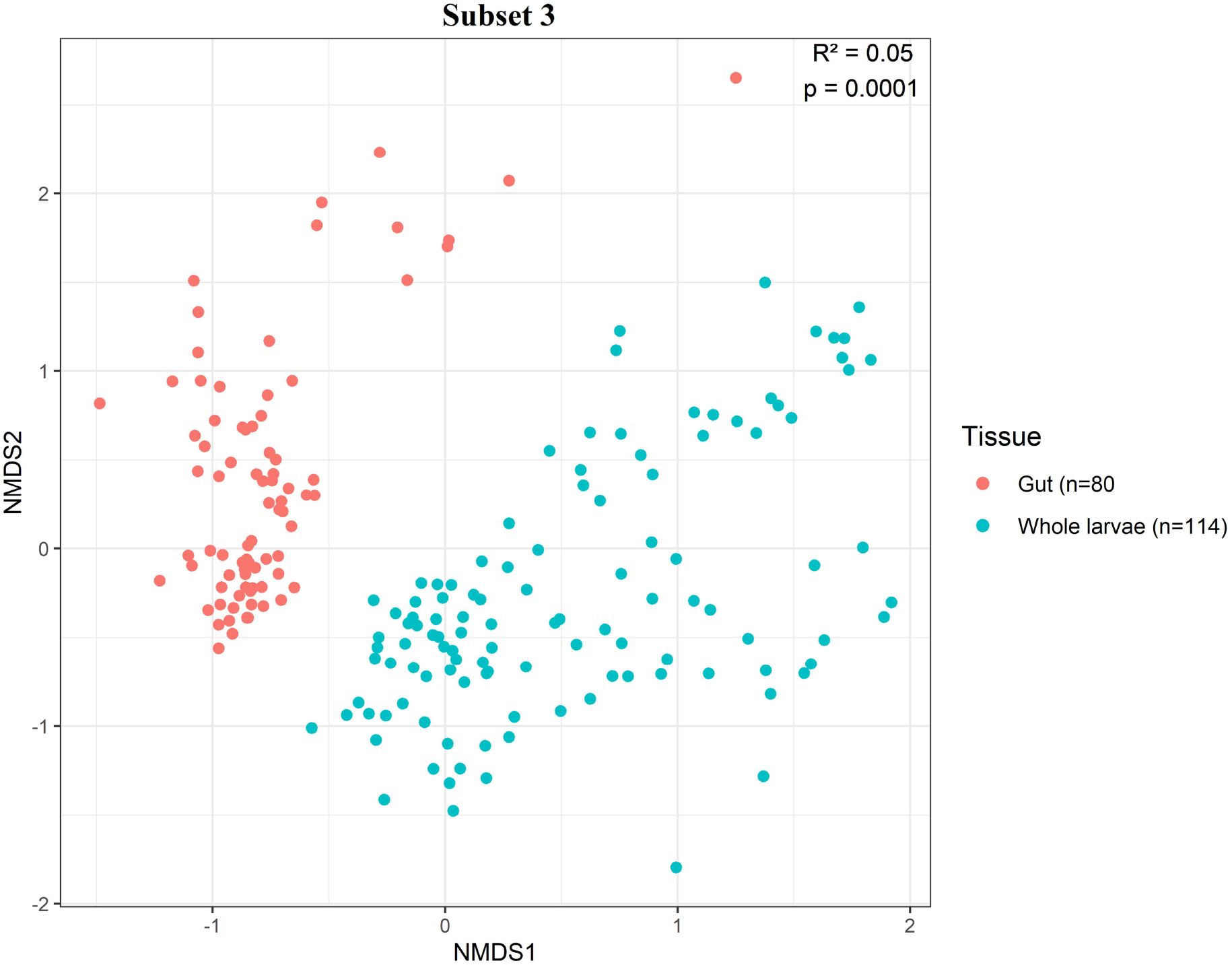
Beta-analysis NMDS plot of Bray-Curtis dissimilarities of Hellinger transformed relative abundance data of Subset 3. Grouping occurred based on the variable factor “Tissue” and distinguishes gut (red, n=80) and whole larvae (blue, n=114) samples. Results of an PERMANOVA analysis are shown in the upper right corner of the plot and reveal the significance (p-value) of the factor “Tissue” and the amount of variation that factor explains (R^2^).

**Table 3.** Results of PERMANOVA analysis of the factor “Tissue” on BSFL microbiota composition. The factor “Tissue” had a significant effect on the microbiota composition found in “Gut” and “Whole larvae” samples within Subsets 3. Statistical testing of Bray-Curtis dissimilarities of Hellinger-transformed relative abundance data was done using PERMANOVA with the adonis function in R with 9,999 permutations.

|  | **Subset 3** |  |
| --- | --- | --- |
| **Factor** | **R<sup>2</sup>** | **P-value** |
| *Tissue* | 0.05 | 0.0001 |

## Discussion

### Core bacterial genera reoccur in BSFL gut samples despite inter-study variations

Despite the variations among the datasets, several bacterial genera showed a high prevalence in both BSFL gut sample subsets. zOTUs representing members of *Enterococcus* and *Morganella* were found in over 80% of the samples and were therefore included in our most strict definition of the core bacteria of BSFL. zOTUs identified as members of the genus *Providencia* were found in more than 75% of all samples in both subsets. The less strict definition of the “core” microbiota also included members of *Scrofimicrobium* and *Klebsiella* in both gut sample subsets. With the exception of *Scrofimicrobium*, members of these genera have also been identified in other BSFL studies which are not included in this meta-analysis (Ao *et al.*, 2020; Gold *et al.*, 2021). *Scrofimicrobium* is a new genus of the *Actinomycetaceae* family, which is closely related to *Actinomyces* and only described so far by Wylensek *et al.* (2020) based on isolates from the pig gut. *Actinomyces* has been found before in BSFL gut studies (Ao *et al.*, 2020; Galassi *et al.*, 2021; Gold *et al.*, 2021). Based on this meta-analysis, members of the above mentioned bacterial genera can be clearly considered as core genera of the BSFL gut community. However, this does not exclude other bacterial genera to belong to the core microbiota of the BSFL gut: increasing the amount of samples in future meta-analyses can provide even stronger insights in omnipresent bacterial genera in the BSFL gut.

Future research can be dedicated to unravel the function(s) of BSFL gut bacteria, as it is still a largely unknown field. For example, *Enterococci* are found in a wide variety of ecological niches, such as in the gastro-intestinal (GI) tracts of human, animal and insect hosts, but also in plants, soil, water and man-made products (Lebreton *et al.*, 2014). Although many *Enterococcus* strains are considered pathogenic (Murray, 1990), Shao *et al.* (2017) found that in the Egyptian cotton leafworm (*Spodoptera littoralis*), the native gut bacterium *Enterococcus mundtii* had an inhibiting effect on other pathogenic enterococcal bacteria such as *Enterococcus faecalis* and on other gram-positive bacteria by altering the environment of the host and thereby increasing host survival (Shoa *et al.*, 2017). *Morganella morganii* has been reported to cause mortality in the Mexican fruit fly (*Anastrepha ludens*) (Salas *et al.*, 2017). The species is, however, also observed to produce phenol, which serves as a sex pheromone in the New Zealand grass grub (*Costelytra zealandica*) (Marshall *et al.*, 2016). The presence of bacteria from the genera *Ignatzschineria* and *Actinetobacter* inhibits oviposition in BSFL, while the presence of certain *Providencia* species promotes this insect behavior (Zheng *et al.*, 2013). *Providencia* has also been proposed to play a role in protein and lipid conversion in the BSFL gut (Ao *et al.*, 2020) and to produce xylanases (Raj *et al.*, 2013), which can assist the insect with the digestion of plant cell walls (Sontowski & van Dam, 2020). However, several members of this genera have also shown various degrees of virulence in fruit flies (*Drosophila melanogaster*) (Galac & Lazzaro, 2011). *Klebsiella*, alongside several other gut bacteria such as *Enterobacter* and *Pseudomonas*, has been reported to produce antioxidants that reduce the amount of toxic oxidants in blood-feeding insects (Sontowski & van Dam, 2020). The presence of these bacteria, that help in digestion or in neutralizing toxic components, might explain why BSFL are able to survive and grow on a wide diversity of substrates. Unravelling the function of BSFL core bacteria can provide more insight in the relation between the insect and its microbiota.

### Experimental parameters affecting the gut bacterial composition

The results of this meta-analysis show that experimental parameters such as larval age at harvest or rearing substrate influences the BSFL gut microbiota composition. This was shown in some earlier studies as well (Ao *et al.*, 2020; Bruno *et al.*, 2019; Osimani *et al.*, 2021; Cifuentes *et al.*, 2020; Zheng *et al.*, 2013), but the added value of a meta-analysis is the possibility to assess whether or to what extent results depend on specific studies. Although some types of substrate were placed in the same category for this analysis, variation between feeds within one category may still be present. For example, chicken feed was often used as a control feed, but the composition of the chicken feed likely varied over different studies. Consequently, its nutritional value and microbial load fluctuated over the studies, which in turn may have been reflected in the microbial composition of the larvae. Similarly, the factor “Age” only differentiated between “Young” and “Old” larvae to make a comparison between grouped studies, yet the microbiota composition of larvae within these subcategories can vary. Because not all studies use the same type of feed and the same larval age at harvest, not all variation can be ascribed solely to these factors. The gut microbiota is more likely influenced by a range of factors and their interactions. Within the parameter “Study”, some factors that potentially influence the microbiota composition might not be captured in this analysis. A relatively large proportion of the microbiota variation is explained by this combination of unknown factors summarized in the factor “Study”, which has also been observed in cattle microbiota studies (Holman & Gzyl, 2019; Holman *et al.*, 2017). One of these factors might be the inhouse bacterial community of rearing facilities used in certain studies. This is best illustrated by the high prevalence of *Dysgonomonas* species in Subset 2, which is made up of two different studies by Klammsteiner *et al.* and one study by Cifuentes *et al.* (Supplementary Figure S4). These four zOTUs associated with *Dysgonomonas* are present in more than 50% of the samples, whereas *Dysgonomonas* is significantly less prevalent in samples from Subset 1. When regarding the number of reads of these four *Dysgonomonas* species, 99.98% of the reads originate from the two studies by Klammsteiner *et al.* and merely 0.02% was found in the study by Cifuentes *et al.* Moreover, a high number of reads of the most prominent *Dysgonomonas* species (zOTU3) was always found in the samples described by Klammsteiner *et al.*, regardless of feed type. This suggests that this species is more likely member of the house flora of the rearing facility, rather than a typical member of the BSFL core species. Other factors such as genetic variation in BSF-strains have not been included in this analysis and it is not known yet whether they should be taken into consideration when evaluating the effect of experimental parameters.

Although core bacteria are found in samples regardless of their experimental treatment, the abundance of these species varied. For example, zOTUs identified as members of the genus *Morganella* in Subset 1 accounted on average for 7.0% of all reads, but the abundance of this genus varied greatly among samples: zOTUs identified as *Morganella* could cover between 0.16% to 95.2% of the reads in a sample (Supplementary Figure S5). This pattern occurred for other core species as well, as the number of reads varied between a minimum of 0-1 % and a maximum of 61.1 % (*Enterococcus*), 75.9% (*Providencia*), 15.8% (*Klebsiella*) and 3.9% (*Scrofimicrobium*). For Subset 2, similar observations can be made (Supplementary Figure S6). These observed variations in number of reads can be attributed to some extent to the specific experimental factors. This can be illustrated by considering the individual samples from the Tegtmeier *et al.* (2021a) dataset as analysed in Subset 1 (Supplementary Figure S5). These BSFL samples all originated from the same insect producer, were used in the same study and were harvested at the same larval instar phase. The main experimental condition remaining in this study was “Feed”, where half of the samples were reared on a control chicken feed and the other half on cottonseed press cake (CPC). zOTUs identified as *Morganella* were present in all samples, but the abundance was significantly higher in the control feed (on average 75.8%) compared to CPC feed (on average 21.7%), indicating that the factor “Feed” influences the relative abundance of *Morganella*. However, not all variation in abundances could be explained by this factor. There was also variation between samples within the same feeding group, where the abundance of *Morganella* was very high in some samples of the CPC group and very low in one of the samples from the control group. This variation in relative abundance, which was also noticeable for other core bacteria, is unlikely related to the experimental conditions (such as relative humidity, rearing conditions, crowding, age and feed), since these were equal for each sample. These examples illustrate the plasticity of the core bacteria: they are prevalent in most samples, but their relative abundance can clearly vary and the exact triggers behind these changes remain to be defined.

### Challenges in performing a meta-analysis of the BSFL microbiota

In any meta-analysis, the aim is to obtain the highest number of similar samples in order to limit the amount of variation and draw general conclusions about the effects of the experimental variables. Data availability and clear sample description is therefore of utmost importance for any meta-analysis. In BSFL next generation sequencing research, there are studies that sequence (part of) the dissected larval gut and studies that use the whole, undissected larvae. In this study, it was shown that there is variation in bacterial species diversity between “Gut” and “Whole larvae” samples. The species bacterial species diversity was higher when whole larvae were examined compared to the larval gut. The higher amount of species diversity observed in the whole larvae might be caused by environmental bacteria that reside on the surface of the insect or enter their cuticula. Future research should carefully consider these variations and involve the correct sample material. Variation in the microbiota composition between different parts of the insect has been demonstrated before. Deguenon *et al.* (2019) found a significant higher bacterial diversity on the external surface of the black blow fly (*Phormia regina*) compared to the gut of the fly. This higher bacterial diversity in external samples has also been found in the house fly (*Musca domestica*) (Park *et al.*, 2019) and in both healthy and parasitized cabbage white (*Pieres brassicae*) caterpillars (Gloder *et al.*, 2021). However, it must be noted that in these studies, the external microbiota was obtained through a washing step of the outer cuticle of the insect and thus these samples did not contain any larval tissue. Nonetheless, due to this kind of variations, it is important to draw separate conclusions regarding the “Gut” microbiota and a more general “Whole Larvae” microbiota. This should be reflected in the terminology used in scientific papers. Research on both the “Gut microbiota” (using dissected samples) and the “Whole microbiota” (using whole larvae) is useful, but each in a different context. Focusing on the “Gut microbiota” is relevant in fundamental studies on the composition and functions of gut micro-organisms. Information on the “Gut microbiota” expands the knowledge on conditions enabling good digestion of (sometimes difficult) substrates (such as some organic waste streams), and hence growth and yield of BSFL production. Considering the “Whole microbiota” is a good approach when studying the microbiological food safety and shelf life of BSFL, other quality aspects in industrial production. Furthermore, since it does not require dissection, analysis of whole larvae samples is less time consuming, which allows for greater sample sizes in these kind of experiments.

In either gut or whole larvae next generation sequencing, the targeted region of the 16S rRNA gene, primer choices and bio-informatical parameters varies among studies. As a result, the data used in this meta-analysis originated from two different regions, with very little overlap between them and resulting in two separate analyses. Variations in 16S rRNA gene regions have been observed by Holman *et al.* (2017), who studied the effect of this parameter on the microbiota composition in the swine gut. When comparing sequencing derived from the V1-V3 and V4 region of the 16S rRNA gene, they found a higher Simpson’s reciprocal value in samples from the V4 region, although there was no variation found in sample richness and Shannon index data (Holman *et al.*, 2017). Holman *et al.* (2017) further noticed that on the genus level some bacteria genera were far less abundant in sequences originating from the V1-V3 region compared to the V4 region. In these cases, the choice of 16S rRNA gene variable region might result in an underestimation of relative abundances of these bacterial genera. In this current study, there were no significant differences in species diversity between gut samples from Subset 1 and Subset 2. Although the results of both analyses are therefore comparable, potential underestimation of bacteria genera in either Subset 1 or 2 due to region selection might be possible. Future meta-analyses would benefit from uniformity among researchers in primer choices and pipeline usage. The usage of universal primers such as (314F (5’-CCTAYGGGRBGCASCAG-3’) and 806R (5’-GGACTACNNGGGTATCTAAT-3’) and limiting the length of used sequences to 250 bp would result in more comparable sequences among studies. Usage of a universal bioinformatics pipeline among microbiota researchers might be unlikely, since some software can be more suitable for specific tasks such as data visualization or statistics and the usage of such programs is often a personal preference of the researcher. Although the analytic methods are often described to some degree, a standardized method of reporting the bioinformatics process could benefit the repeatability of these bioinformatic analyses. This includes information about used software, sequence length and estimate error thresholds, methods of data filtering and referenced databanks, which all could be added in a supplementary table.

Variation in data generation is not the only pitfall of BSFL microbiota analysis. Interpretation of results can be influenced by the use of alternative definitions of the “core “ microbiota. In this study, the intrinsic core bacterial community of the BSFL was defined as those bacterial genera that were found above a prevalence threshold of either 80% or 50% across multiple samples, regardless of external factors (*i.e.* experimental setup, larval age, rearing facility). This definition follows the same criteria as the “Common core” definition by Risely (2020) and it can be quantified by scoring the presence or absence of bacteria found in samples, as described by Shade & Handelsman (2012). This method does not take into account the relative abundance of bacterial taxa found in the samples (Shade & Handelsman, 2012), nor does it provide information about the function of the bacterial species or their dynamics over time (Risely, 2020). The definition of “core” bacterial species varies greatly among (black soldier fly) studies: the core microbiota can be described as bacterial groups present in all samples regardless of environmental and/or experimental conditions (Ao *et al.*, 2020), as groups being abundant in a certain percentage of samples (Klammsteiner *et al.*, 2020; Shelomi *et al.*, 2020) or as groups persistent over life stages and time (Cifuentes *et al.*, 2020; Raimondi *et al.* 2020; Zhan *et al.*, 2020). Several definitions can exist next to each other, as long as studies clearly describe the definition(s) they consider, so that comparisons between studies can be made correctly.

## Conclusions

This study evaluated the effects of experimental parameters and bio-informatical choices on the bacterial composition of BSFL, using data originating from dissected guts as well as whole larvae. This approach revealed variation in species diversity between the sample types. Hence, including only the gut or the whole larvae is important to consider when designing future BSFL microbiota experiments, and the choice should be aligned with the aim of the study. Through the meta-analysis, core gut bacterial genera were identified. Regardless of encountered variations, members of the genera *Enterococcus, Morganella* and *Providencia* were the most prevalent in both subsets of BSFL gut samples, followed by *Klebsiella* and *Scrofimicrobium*. Furthermore, there is room for improvement in standardizing DNA extraction methods and 16S rRNA gene sequence processing, so that more BSFL sequencing data can be combined and analysed in future research. This will allow more profound insights and eventually practical recommendations for BSFL production.

## Funding

This work was supported by the Research Foundation Flanders (FWO Vlaanderen) [grant numbers S008519N, 12V5219N].

## Conflict of interest

The authors declare that there are no conflicts of interest.

## Legends supplementary figures

**Supplementary Figure S1. Rarefaction curves of Subset 1.** Colors indicate from which study the sample originates from.

**Supplementary Figure S2. Rarefaction curves of Subset 2.** Colors indicate from which study the sample originates from.

**Supplementary Figure S3. Rarefaction curves of Subset 3.** Colors indicate from which study the sample originates from.

**Supplementary Figure S4. Total number of reads from four zOTUs from Subset 2, all identified as members of the genus *Dysgonomonas.*** X-axis shows the feed types used in these three separate studies, y-axis show the total number of reads for each zOTU. In the Cifuentes *et al.* dataset, only zOTU8 was found with a total of 206 reads.

**Supplementary Figure S5. Overview of all samples and their relative abundance of the core zOTUs of Subset 1.** Reads from zOTUs other then the core genera are clustered together as “Other”. Highlighted are the samples from the study of Tegtmeier *et al.* (2021a), starting with codes SRR1300. Red encircled samples were reared on a cottonseed press cake substrate, green encircled samples were reared on a control chicken diet.

**Supplementary Figure S6. Overview of all samples and their relative abundance of the 20 core zOTUs of Subset 2.** Reads from zOTUs other then the core species are clustered together as “Other”.

## Notes

### Competing Interest Statement

The authors have declared no competing interest.

