## Supplementary figures and images for "Meta-analysis of the black soldier fly (*Hermetia illucens*) microbiota based on 16S rRNA gene amplicon sequencing"

### Supplementary Figure 1

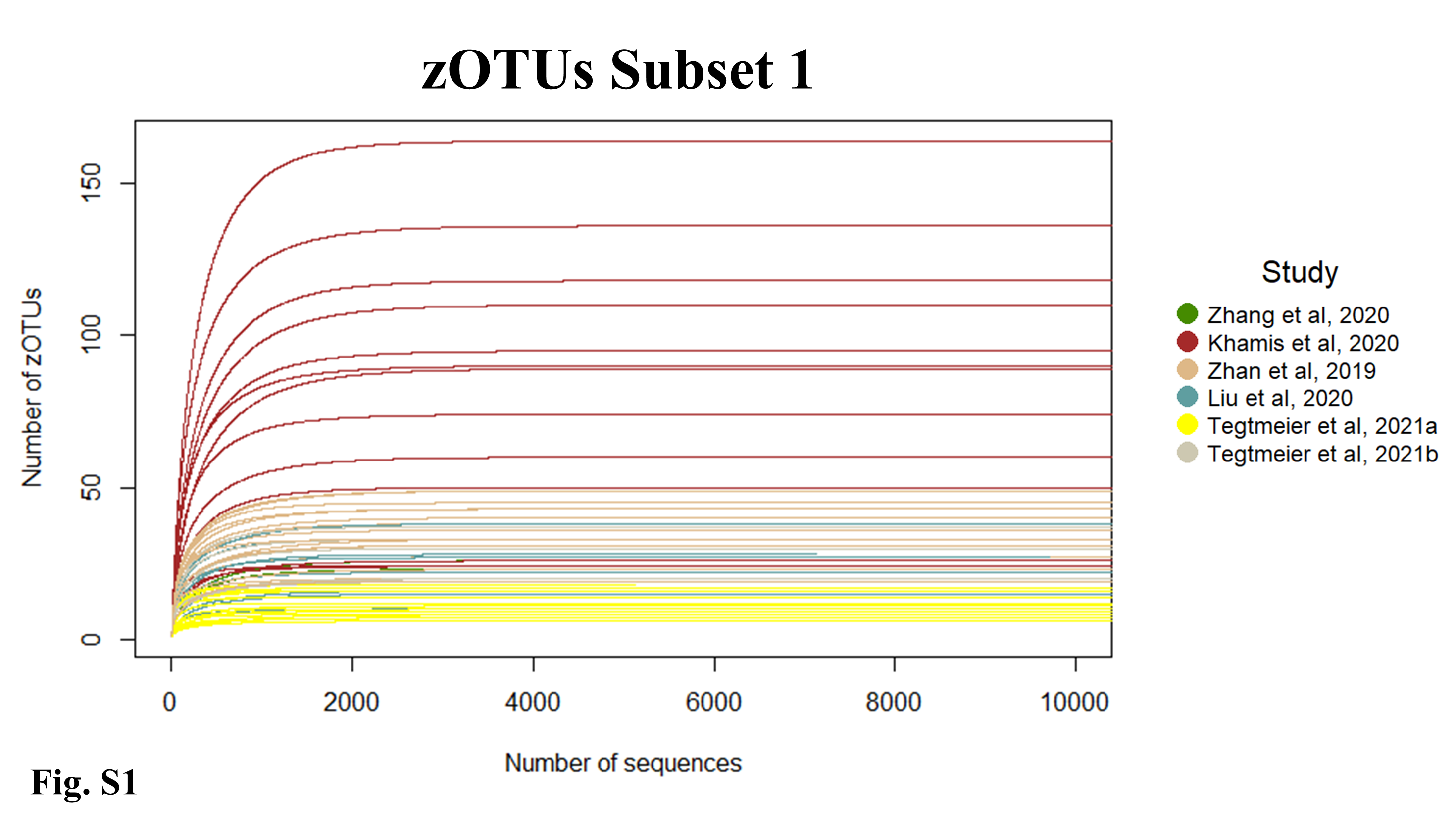

### Supplementary Figure 2

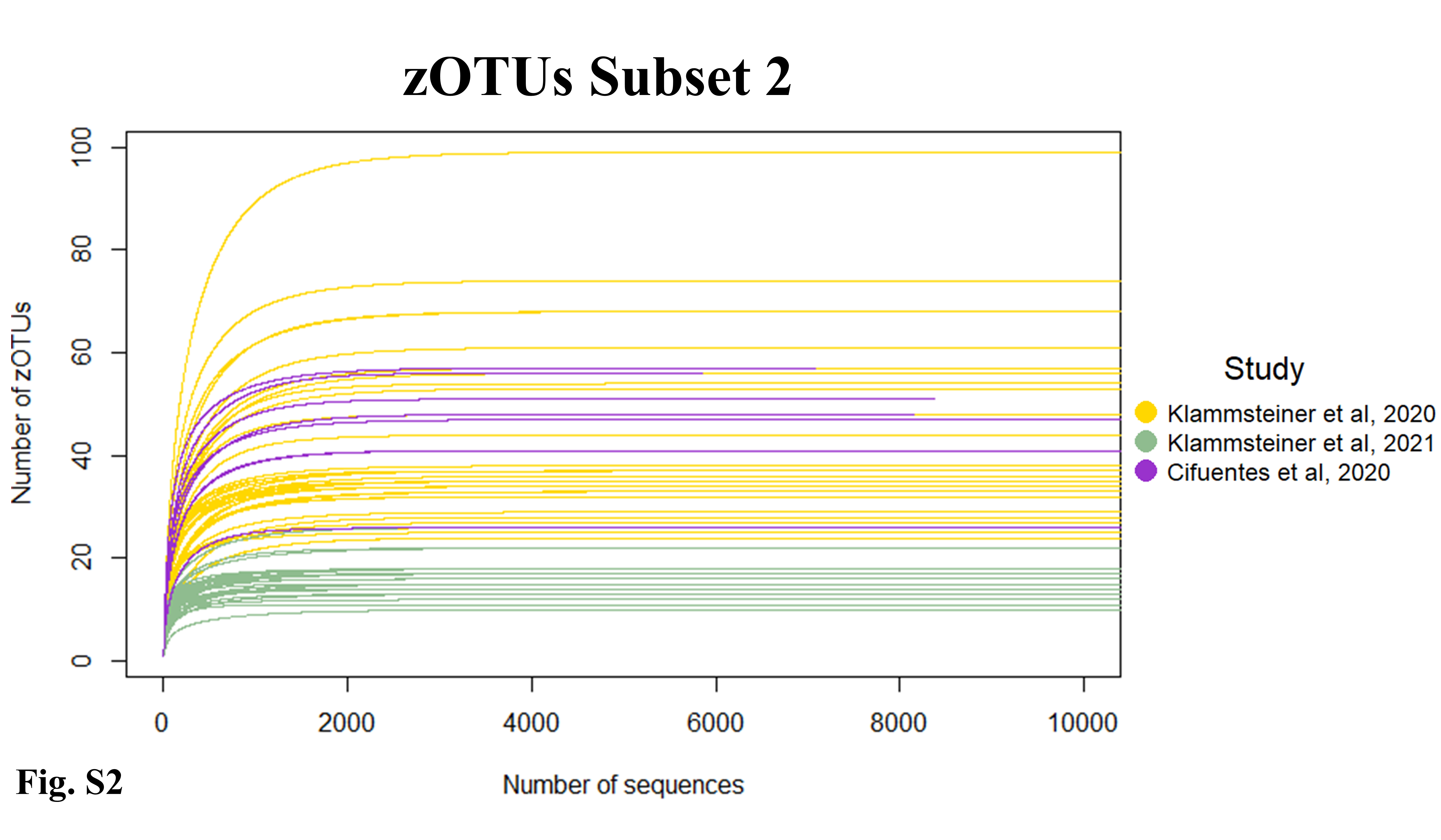

### Supplementary Figure 3

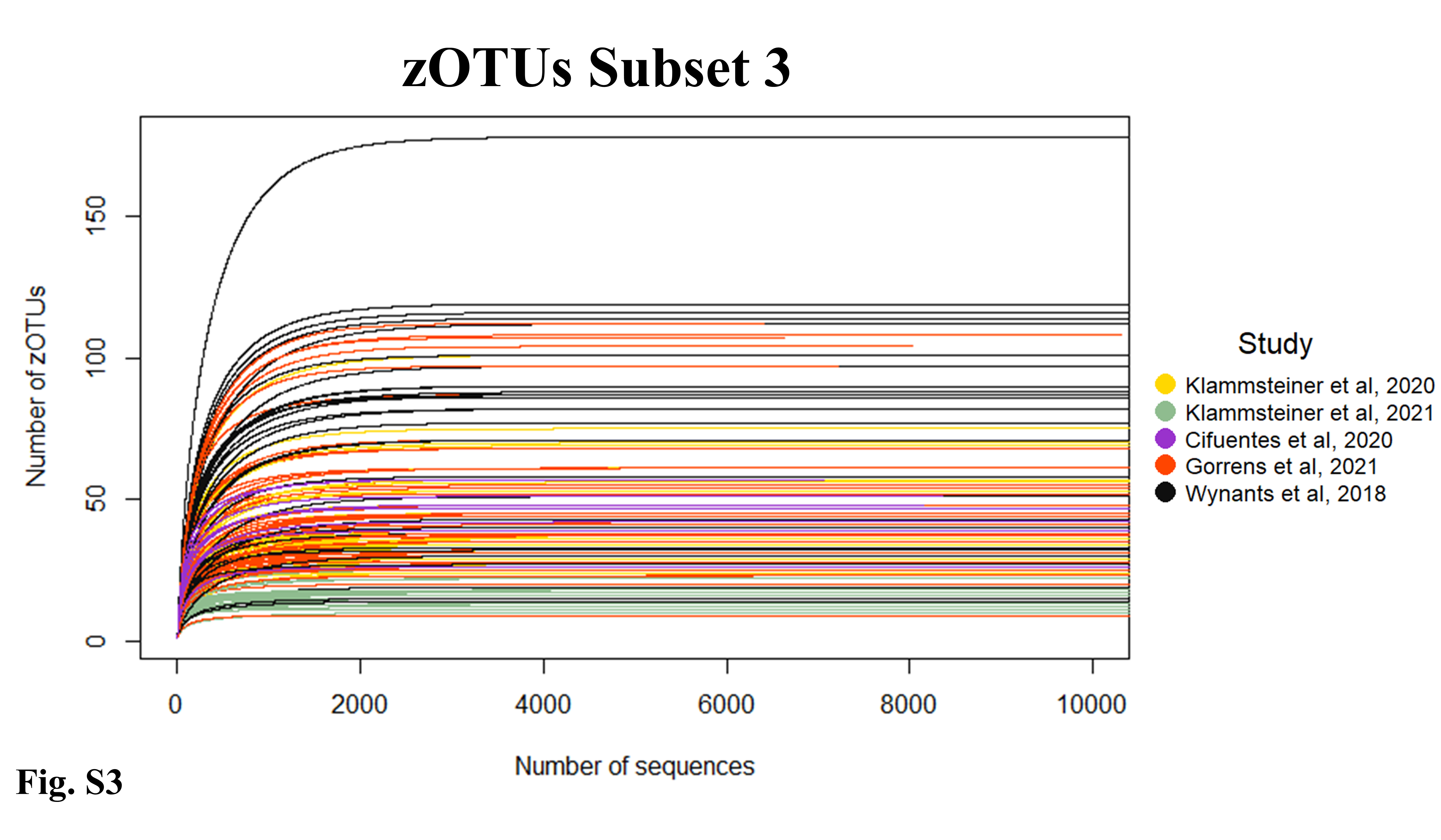

### Supplementary Figure 4

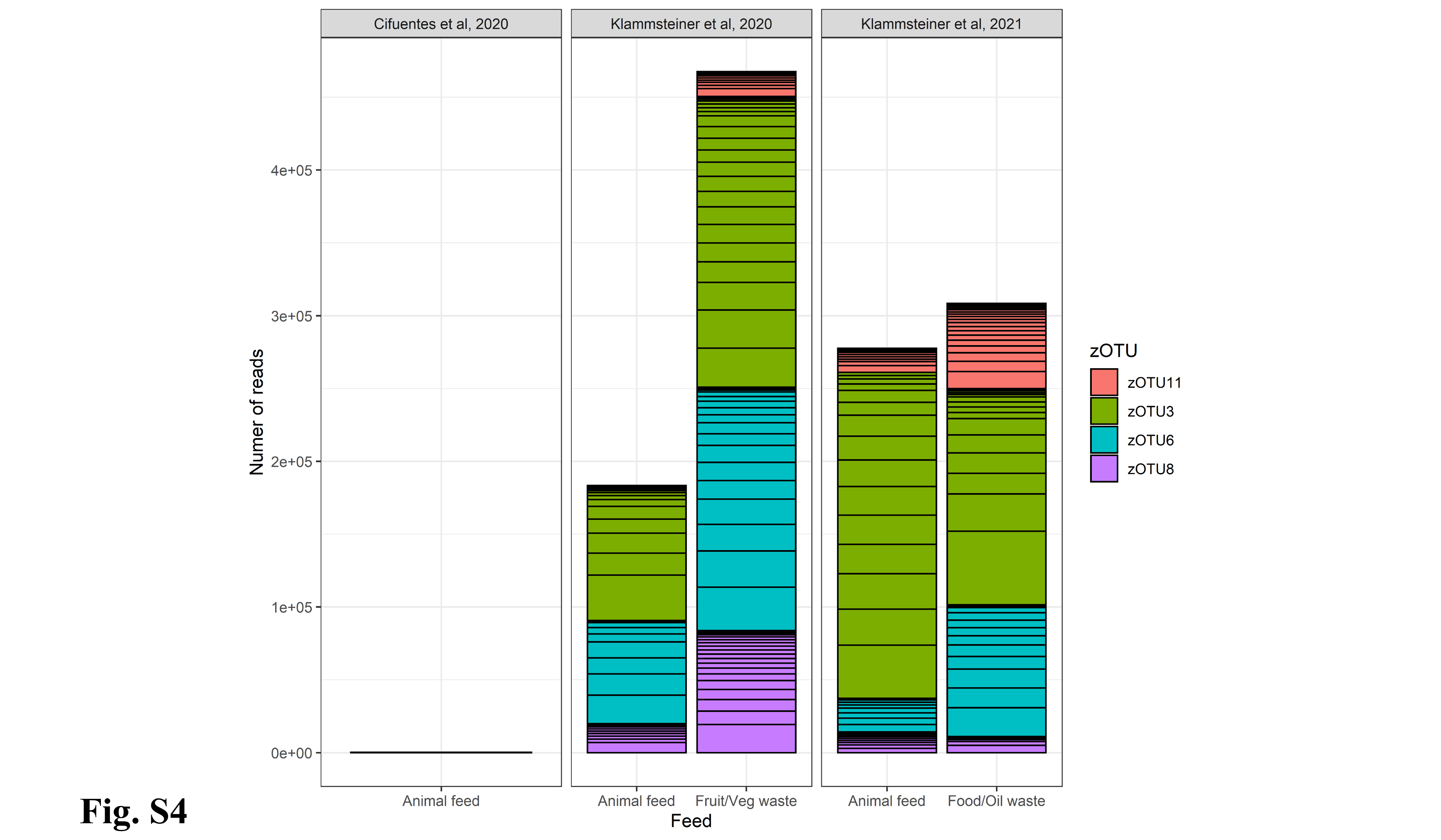

### Supplementary Figure 5

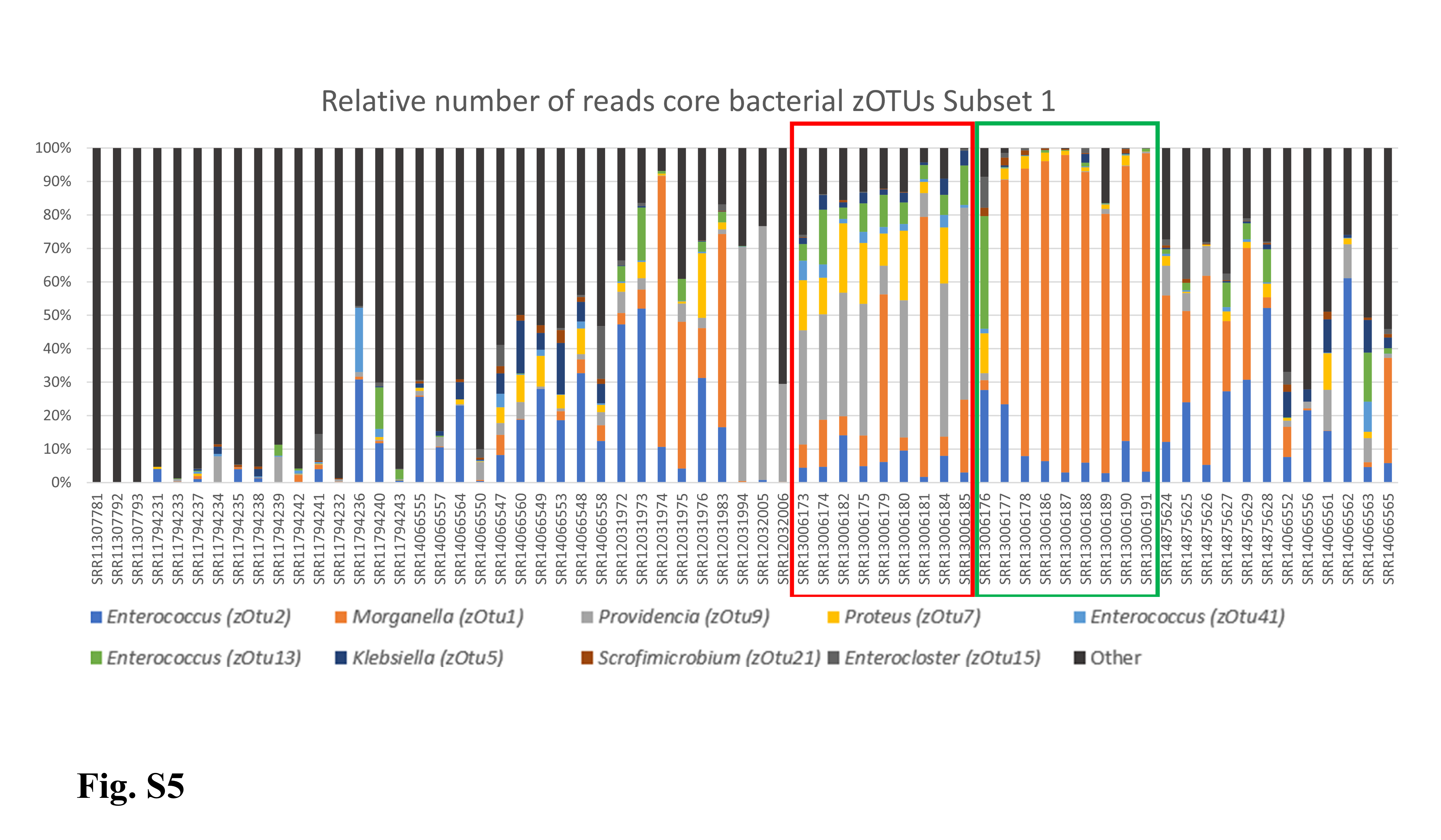

### Supplementary Figure 6

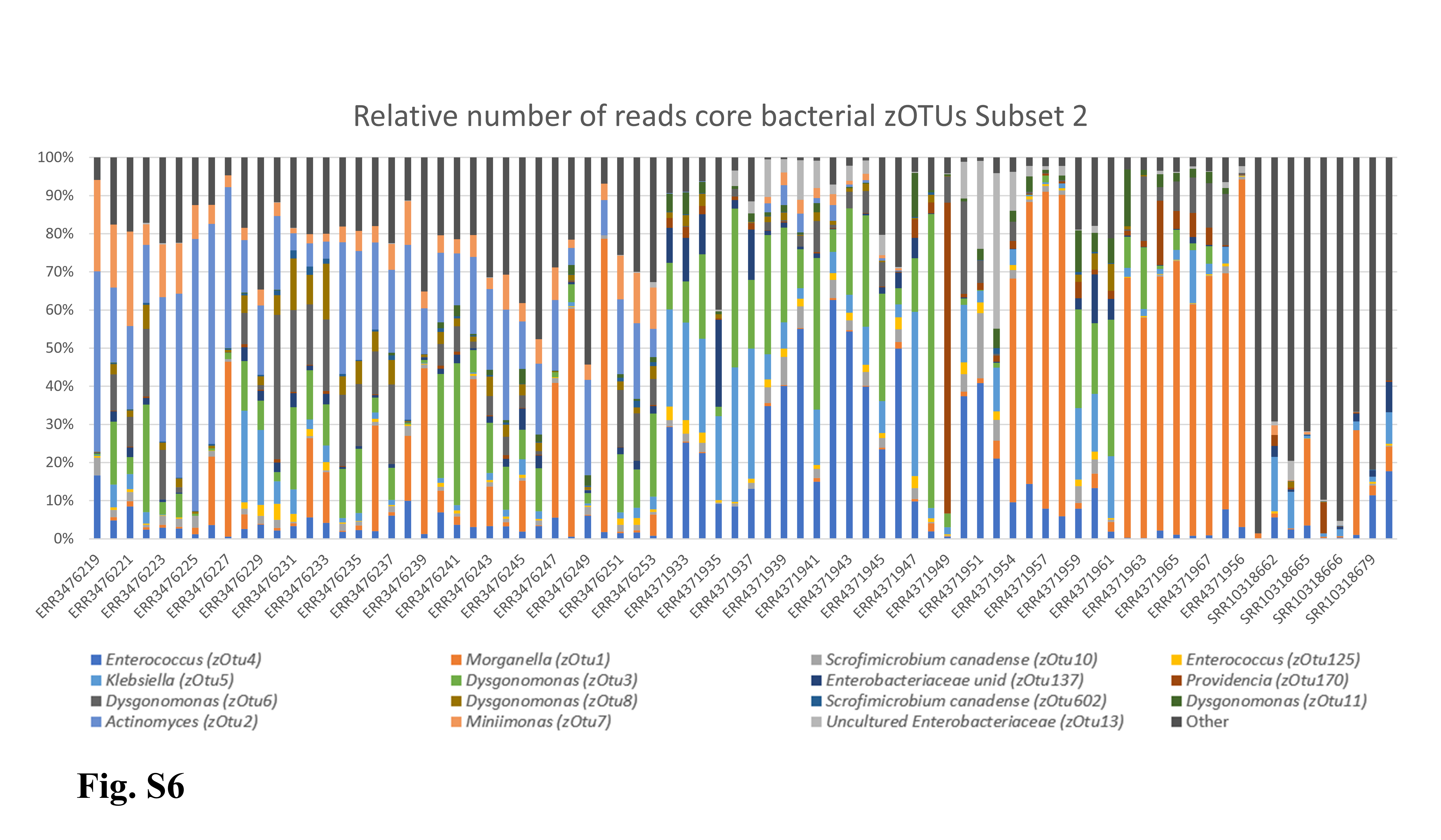
