## Supplementary Tables for "Meta-analysis of the black soldier fly (*Hermetia illucens*) microbiota based on 16S rRNA gene amplicon sequencing"

**Supplementary Table S1.** List of BSFL microbiota studies that were considered but due to varying reasons could not be included in this meta-analysis.

| Study | Reason for rejection | Reference |
| --- | --- | --- |
| Bruno *et al.,* 2019 | Mismatch in sample type: differentiation between regions of the midgut out of the scope of this study | https://doi.org/10.1128/AEM.01864-18 |
| Cai *et al.*, 2018a | Raw sequence data not available | https://doi.org/10.1111/1462-2920.14450 |
| Cai 2018b | Raw sequence data not available | https://doi.org/10.1016/j.envpol.2018.06.105 |
| Galassi *et al.*, 2021 | Raw sequence data not available | https://doi.org/10.1093/jisesa/ieaa148 |
| Gold *et al.*, 2021 | No separate forward and reverse available from datasbase. Merged sequences have too high expected number of errors at a length of 250 bp | https://doi.org/10.3920/JIFF2021.0038 |
| Huang *et al.*, 2020 | Experiments were conducted on genetically modified and sterilized BSFL. Feeds were artificially inoculated with bacteria. Samples were outside of the scope of this meta-analysis. | https://doi.org/10.1371/journal.pone.0225873 |
| Jiang *et al.*, 2019 | No separate forward and reverse available from datasbase. Merged sequences have too high expected number of errors at a length of 250 bp | https://doi.org/10.1111/1751-7915.13393 |
| Kooienga *et al.*, 2020 | Metadata did not provide enough information to differentiate treatment among samples | https://doi.org/10.3389/fmicb.2020.587979 |
| Osimani *et al.*, 2020 | Only study providing whole larvae on the group 1 region, no comparison with other whole larvae samples. Furthermore, artificial enriched substrates outside of the scope of this research | https://doi.org/10.1016/j.foodres.2020.110028 |
| Raimondi *et al.*, 2020 | Sequences of length of the samples too short (190 bp) | https://doi.org/10.3390/microorganisms8060902 |
| Schreven *et al.*, 2021 | Quality score of available raw sequences to low to allowing further processing of the data | https://doi.org/10.1093/femsec/fiab054 |
| Shelomi *et al.*, 2020 | No separate forward and reverse available from datasbase. Merged sequences have too high expected number of errors at a length of 250 bp | https://doi.org/10.1093/ee/nvz164 |
| Tanga *et al.*, 2021 | Raw sequence data not available | https://doi.org/10.3389/fmicb.2021.635881 |
| Wu *et al.*, 2020 | Raw sequence data not available | https://doi.org/10.1016/j.ecoenv.2020.110323 |
| Yang *et al.*, 2021 | Raw sequence data not available | https://doi.org/10.3389/fmicb.2021.601253 |
| Zhan *et al.*, 2020 | Raw sequence data not available | https://doi.org/10.1038/s41422-019-0252-6 |
| Zhineng *et al.*, 2021 | Raw sequence data not available | https://doi.org/10.1186/s13213-021-01626-8 |

**Supplementary Table S2.** Overview of the top 20 most prevalent zOTUs found in Subset 1 (left) and Subset 2 (right), based on an absence/presence criterium. BLAST was used to identify the zOTUs. Names of the matching bacterial genera as well as the accession number and E-value are shown. Furthermore, the relative abundance of each zOTU across the whole group is given.

| zOTUs Subset 1 | Nucleotide collection (nt) % identification | Accession number nt database | E-value | Prevalence (% total samples) | Abundance (% total samples) | zOTUs Subset 2 | Nucleotide collection (nt) % identification | Accession number nt database | E-value | Prevalence (% total samples) | Abundance (% total samples) |
| --- | --- | --- | --- | --- | --- | --- | --- | --- | --- | --- | --- |
| zOTU2 | *Enterococcus* (93.50%) | KX523839.1 | 6e-96 | 87.7% | 15.0% | **zOTU4** | *Enterococcus* (99.60%) | NR_159231.1 | 8e-126 | 98.8% | 10.4% |
| zOTU1 | *Morganella* (100%) | NR_113580.1 | 2e-127 | 84.6% | 7.0% | **zOTU1** | *Morganella* (100%) | LS992078.1 | 2e-127 | 95.0% | 19.2% |
| zOTU9 | *Providencia* (100%) | MK503720.1 | 2e-127 | 80.0% | 3.9% | **zOTU10** | *Scrofimicrobium canadense* (98.01%) | MN537492.1 | 4e-119 | 90.0% | 1.9% |
| zOTU7 | *Proteus* (100%) | MN749810.1 | 2e-127 | 73.8% | 2.6% | **zOTU125** | *Enterococcus* (99.60%) | NR_159231.1 | 8e-126 | 90.0% | 1.1% |
| zOTU41 | *Enterococcus* (100%) | LC521987.1 | 2e-127 | 69.2% | 1.3% | **zOTU5** | *Klebsiella* (100%) | MN272345.1 | 2e-127 | 90.0% | 6.9% |
| zOTU13 | *Klebsiella* (100%) | NR_114152.1 | 2e-127 | 64.6% | 1.7% | **zOTU3** | *Dysgonomonas* (97.20%) | MT340883.1 | 3e-114 | 82.5% | 11.5% |
| zOTU5 | *Enterococcus* (99.18%) | NR_159231.1 | 8e-121 | 64.6% | 3.9% | **zOTU137** | Unidentified *Enterobacteriaceae* (99.20%) | MT256272.1 | 4e-124 | 80.0% | 2.3% |
| zOTU21 | *Scrofimicrobium canadense* (99.18%) | MN537492.1 | 8e-121 | 56.9% | 0.9% | **zOTU170** | *Providencia* (100%) | MK503720.1 | 4e-124 | 75.0% | 2.8% |
| zOTU15 | *Enterocloster* (97.96%) | NR_042200.1 | 9e-116 | 52.3% | 1.5% | **zOTU6** | *Dysgonomonas* (100%) | NR_113134.1 | 9e-111 | 73.8% | 6.1% |
| zOTU72 | *Klebsiella* (100%) | MN894279.1 | 8e-126 | 38.5% | 0.5% | **zOTU8** | *Dysgonomonas* (100%) | NR_113133.1 | 2e-127 | 71.3% | 2.2% |
| zOTU6 | *Ignatzschineria* (99.18%) | LC377581.1 | 8e-121 | 38.5% | 6.6% | **zOTU602** | *Scrofimicrobium canadense* (99.20%) | MN537492.1 | 4e-124 | 63.8% | 0.4% |
| zOTU3475 | *Enterococcus* (99.20%) | LT576388.1 | 4e-124 | 38.5% | 0.2% | **zOTU11** | *Dysgonomonas* (100%) | NR_137388.1 | 2e-117 | 63.8% | 1.6% |
| zOTU8 | *Proteus* (100%) | MN749810.1 | 2e-127 | 35.4% | 1.1% | **zOTU2** | *Actinomyces* (97.60%) | LT855383.1 | 2e-117 | 58.8% | 11.6% |
| zOTU73 | *Streptomyces* (95.14%) | MH144589.1 | 1e-104 | 33.8% | 0.1% | **zOTU7** | *Miniimonas* (98.40%) | NR_112996.1 | 8e-121 | 58.8% | 3.7% |
| zOTU4 | *Romboutsia* (99.18%) | NR_144740.1 | 8e-121 | 32.3% | 7.9% | **zOTU13** | Uncultured *Enterobacteriaceae* (97.60%) | JN423140.1 | 2e-115 | 57.5% | 1.9% |
| zOTU37 | *Actinomyces* (97.08%) | LT855383.1 | 3e-110 | 30.8% | 0.7% | **zOTU16** | *Fusicatenibacter* (100%) | MT269542.1 | 6e-126 | 47.5% | 0.7% |
| zOTU26 | *Schaalia* (93.95%) | NR_025366.1 | 4e-99 | 29.2% | 0.8% | **zOTU22** | *Brucella* (99.60%) | NR_146362.2 | 8e-126 | 43.8% | 0.5% |
| zOTU12 | *Clostridium* (99.19%) | AB971795.1 | 7e-122 | 27.7% | 2.7% | **zOTU100** | *Dysgonomonas* (98.80%) | NR_137388.1 | 2e-122 | 42.5% | 1.3% |
| zOTU29 | *Dysgonomonas* (96.73%) | KR822451.1 | 8e-110 | 26.2% | 0.7% | **zOTU18** | *Lacrimispor*a (100%) | AB971794.1 | 2e-127 | 42.5% | 0.6% |
| zOTU44 | *Citrobacter* (100%) | MN548424.1 | 2e-127 | 24.6% | 0.2% | **zOTU17** | *Escherichia/Shigella* (100%) | MN307293.1/ MK184303.1 | 2e-127 | 40.0% | 0.8% |
